# Universal Spectrum Identifier for mass spectra

**DOI:** 10.1101/2020.12.07.415539

**Authors:** Eric W. Deutsch, Yasset Perez-Riverol, Jeremy Carver, Shin Kawano, Luis Mendoza, Tim Van Den Bossche, Ralf Gabriels, Pierre-Alain Binz, Benjamin Pullman, Zhi Sun, Jim Shofstahl, Wout Bittremieux, Tytus D. Mak, Joshua Klein, Yunping Zhu, Henry Lam, Juan Antonio Vizcaíno, Nuno Bandeira

**Affiliations:** Institute for Systems Biology, 401 Terry Ave N, Seattle, WA, 98109, USA; European Molecular Biology Laboratory, European Bioinformatics Institute (EMBL-EBI), Wellcome Trust Genome Campus, Hinxton, Cambridge CB10 1SD, United Kingdom; Center for Computational Mass Spectrometry, Department of Computer Science and Engineering, Skaggs School of Pharmacy and Pharmaceutical Sciences, University of California, San Diego, 92093-0404, USA; Toyama University of International Studies, 930-1292 Toyama, Japan; VIB - UGent Center for Medical Biotechnology, VIB, Ghent, Belgium; Department of Biomolecular Medicine, Faculty of Medicine and Health Sciences, Ghent University, Ghent, Belgium; Clinical Chemistry Service, Lausanne University, 1011 Lausanne, Switzerland; Thermo Fisher Scientific, 355 River Oaks Parkway San Jose, CA 95134, USA; Skaggs School of Pharmacy and Pharmaceutical Sciences, University of California San Diego, La Jolla, CA 92093, USA; Department of Computer Science, University of Antwerp, 2020 Antwerp, Belgium; Mass Spectrometry Data Center, National Institute of Standards and Technology, 100 Bureau Drive, Gaithersburg, MD 20899, USA; Program for Bioinformatics, Boston University, Boston, MA 02215, USA; National Center for Protein Sciences (Beijing), Beijing Institute of Lifeomics, #38, Life Science Park, Changping District, Beijing 102206, China; Department of Chemical and Biological Engineering, the Hong Kong University of Science and Technology, Clear Water Bay, Hong Kong

**Keywords:** Proteomics Standards Initiative, PSI, mass spectrometry, proteomics, Universal Spectrum Identifier, USI, standards

## Abstract

Mass spectra provide the ultimate evidence for supporting the findings of mass spectrometry (MS) proteomics studies in publications, and it is therefore crucial to be able to trace the conclusions back to the spectra. The Universal Spectrum Identifier (USI) provides a standardized mechanism for encoding a virtual path to any mass spectrum contained in datasets deposited to public proteomics repositories. USIs enable greater transparency for providing spectral evidence in support of key findings in publications, with more than 1 billion USI identifications from over 3 billion spectra already available through ProteomeXchange repositories.

## Introduction

Mass spectrometry (MS) is a major analytical tool for the fields of proteomics and metabolomics among others, enabling high throughput identification and quantitation of millions of different analytes including e.g. peptides and small molecules. The availability of proteomics data in the public domain has increased dramatically in recent years due to the wide adoption of open data practices in the field, which has been enabled by the reliability of the public resources that are part of the ProteomeXchange^1^ (PX) consortium. But while the public availability of the raw mass spectrometry data creates the possibility for scientists to directly inspect whether the data support published results as claimed in the corresponding papers, this remains difficult to implement in practice because there is no standard mechanism supported by the repositories to directly reference individual mass spectra whose interpretations are crucial to the published results. As it stands today, the key spectral evidence for biological studies lacks ‘FAIRness’ (i.e. Findable, Accessible, Interoperable and Reusable)^2^ and is at best found as a figure in the manuscript or in the supplementary information, which is typically of low quality and does not provide the connection to the raw data (as is required for inspection of the results for the correctness of the corresponding claims). This need becomes even more pressing as the promise of open data for the advancement of science begins to be realized by data reanalyses revealing multiple cases where the data do not support published results^3,4^. We thus propose a new standardized mechanism supported by proteomics repositories for referring to specific spectra of high importance, suitable for citation of data in published manuscripts, as well as for supporting both human interactions and automated access to the data via application programming interfaces (APIs).

A related mechanism for identifying identical metabolomics mass spectra across repositories has been proposed as the SPLASH identifier^5^, an algorithm-generated hash of the spectrum data. SPLASH was designed to determine if the same exact reference spectrum is present in multiple metabolomics resources, since an identical spectrum generates an identical SPLASH identifier. However, (i) SPLASH was based on statistics of metabolomics spectra that do not apply to proteomics, (ii) requires pre-computing billions of hashes in advance to make them findable and (iii) uses hashes that are not interpretable by humans, and thus is not suitable as a general-purpose identifier for all spectra in proteomics data sets deposited to public repositories. Here we present an alternative approach using a concatenated multi-part key that specifies the collection, MS run, and index information needed to locate a particular mass spectrum in a repository, defining keys that researchers can easily compose without requiring any special hashing algorithms. This concept has been developed by the Human Proteome Organization^6^ (HUPO) Proteomics Standards Initiative^7^ (PSI) as the Universal Spectrum Identifier (USI). It provides a standardized mechanism for identifying a mass spectrum via an abstract path to its location that can be easily mapped to a local path at each resource that contains the spectrum.

The full specification document describing in detail all aspects of version 1.0 of this standard as well as future updates to this specification and information about current implementations with examples are available on the USI page at the PSI web site at http://psidev.info/usi. In this letter, we provide a brief overview of the USI’s components and describe prominent current applications.

## Description of the identifier

The USI is a multipart key separated by colon characters. The basic form of the USI has five components with an optional sixth as depicted in Figure 1. The first component is the prefix “mzspec” to make clear that the following string is a USI; mzspec is a registered namespace at identifiers.org^8^ to aid in automated mapping of a USI string to a URL (https://registry.identifiers.org/registry/mzspec). The second component is the collection identifier, initially intended to be a ProteomeXchange dataset (PXD) identifier^9^, but can be extended to other types of collection identifiers, and can take on the special value of USI000000 as a placeholder for subsequent dataset submission. The third component is the MS run name, typically the root name of the raw vendor format file or mzML^10^ file, either with or without the file extension, depending on the use case as fully described in the specification. The fourth component is a type flag to indicate how to interpret the fifth component; currently allowed values are “scan”, “index”, or “nativeId” (as defined in the mzML specification available at http://psidev.info/mzML). The fifth component is the index value (based on the flag type) within the specified MS run. While these five components can define the abstract path to a spectrum, one additional (sixth) component can specify an interpretation of the spectrum when one is known or proposed. There are many nuances to the specification and the usage of these components that are described in detail in the full specification document.

**Figure 1:**
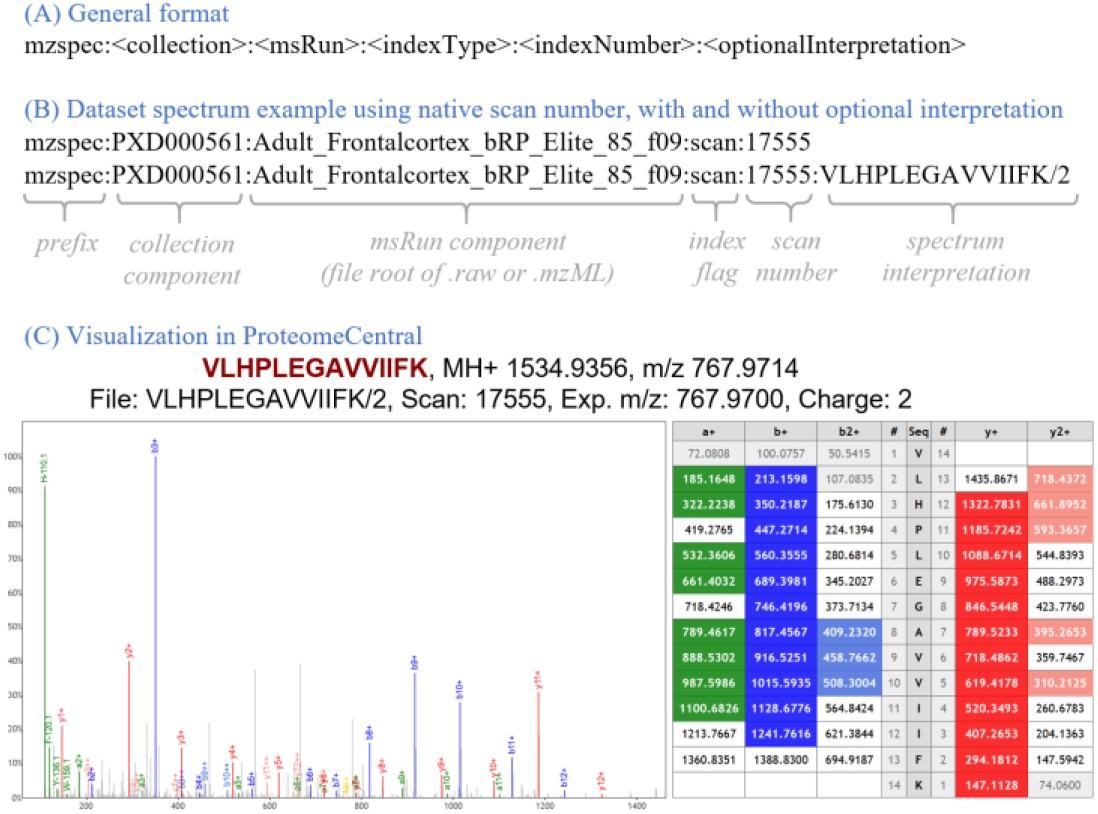
(A) Graphical overview of the general format of the Universal Spectrum Identifier, including the mzspec prefix, the collection component, the MS run component, the indexType, the indexNumber, and the optional interpretation. (B) A USI example using the spectrum scan number, once with and once without the optional spectrum interpretation. (C) Visual representation of this spectrum (mzspec:PXD000561:Adult_ Frontalcortex_bRP_Elite_85_f09:scan:17555:VLHPLEGAVVIIFK/2) in ProteomeCentral, accompanied with the ion table indicating the m/z values of the identified b- and y-ions. The Lorikeet spectrum viewer is used.

The spectrum interpretation, depicted as a simple peptide sequence and charge in Figure 1, can become quite complex when taking into account different types of mass modifications, adducts, and non-peptidic molecules, among other scenarios. An entire separate PSI specification, called ProForma 2.0, has been developed to define the rules for representing complex peptidoforms and proteoforms in a standardized manner. The current version is available at the PSI web page http://psidev.info/proforma.

In addition to the basic form described above, there are several derivative forms that can be used as identifiers of related concepts. If only the first three components are provided, this naturally forms an MS run identifier, or, if a file extension is provided, a particular version of that MS run.

## Applications

There are already several implementations of the USI, and more are emerging. At the spectrum viewer applications of the PX resources PRIDE^11^, PeptideAtlas^12^, MassIVE^13^, jPOST^14^, and iProX^15^, the USI is displayed for each spectrum viewed when the data set containing the spectrum is associated with a PXD identifier, and these resources also provide a text box in which a USI can be pasted in order to display the requested spectrum (see Supplementary Table 1 for listing of URLs, and Supporting Information for a description of the implementations at each site). For convenience, the ProteomeCentral resource of PX implements a single endpoint at http://proteomecentral.proteomexchange.org/usi/ that reaches out to all participating partners to fetch spectra for a provided USI if available at any resource. It also provides USI validator functionality, which checks that the format of the USI complies with the USI specification. An overview of the USI ecosystem is depicted in Figure 2.

**Figure 2.**
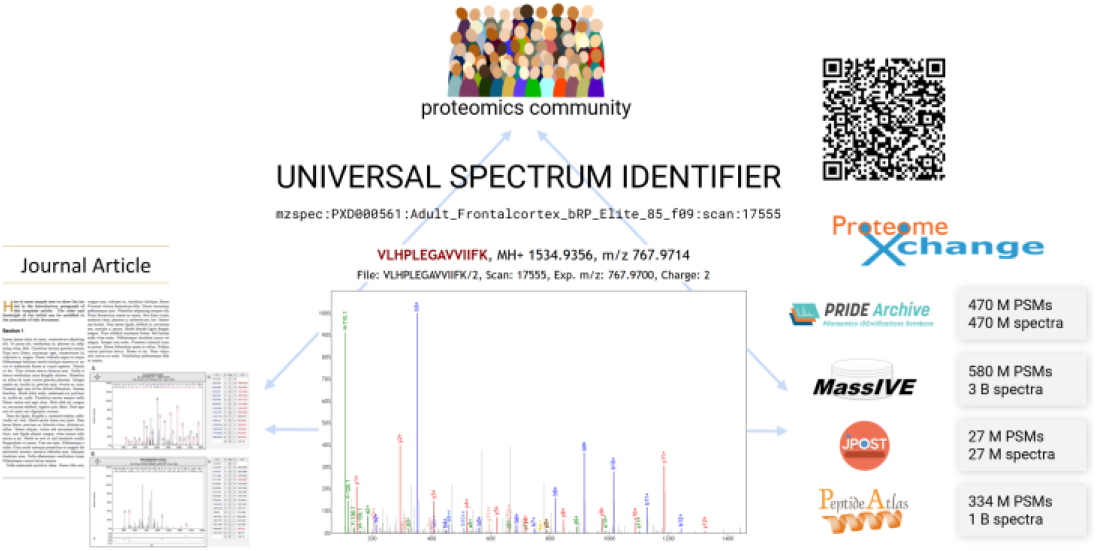
Graphical depiction of USI application ecosystem. Members of the community can uniquely identify spectra from journal articles and other sources using USIs. A USI can be resolved potentially at any of several different repositories that store datasets, or spectra can be obtained and viewed using independent applications, such as at ProteomeCentral, which store no spectra themselves, but can fetch spectral data from repositories using USIs. Hundreds of millions of peptide-spectrum matches (PSMs) and spectra without matches are accessible via USIs at the various repositories. PSMs or spectra can even be uniquely identified with QR codes.

Since the USI is designed as a virtual path to a spectrum based on a set of rules and conventions, there is no need to compute hashes for billions of existent spectra or newly generated spectra like in the case of the SPLASH identifier. Implementing repositories need only map the virtual path components (collection, MS run, indexType) to their local storage system to make the spectra findable and accessible. As a key point, USIs can be retroactively determined for spectra deposited long ago to a ProteomeXchange repository (the spectrum depicted in Figure 1 was deposited in 2013). Furthermore, authors of new publications can easily construct and provide within their articles references to key spectra that support their conclusions by knowing the collection identifier (typically known as a manuscript is being submitted), the MS run filename, and spectrum scan number, which are usually determinable from within most analysis environments. ProteomeXchange repositories also provide various tools to assist with the process of generating USIs for dataset spectra (see Supplementary Materials). Within some proteomics analysis workflows, the provenance information about the original scan numbers is unfortunately lost during processing, but the importance of provenance is now well understood and any such loss of provenance should be remedied in future software revisions. Also, in some cases, deposited spectra may not be accessible because they are stored in a format that is not readable by the hosting resource’s software (for example if the spectra were only deposited as native vendor binary files that have not been converted to open formats), or represent consensus spectra, demultiplexed DIA (Data Independent Acquisition) spectra, or other challenging scenarios that are discussed in detail in section 3.5 of the full USI specification.

In Box 1 we provide a set of 13 example USIs along with a brief comment on each. These same 13 USIs can be easily viewed as the “Box 1 example USIs” select list at http://proteomecentral.proteomexchange.org/usi. Example 4c in Box 1 provides the USI for the demonstrated correct PSM of an ordinary UniProtKB protein Q9UQ35 from Mylonas *et al*.^4^ Figure 2B (example 4d is the corresponding synthetic peptide spectrum). Example 4a in Box 1 provides the USI for the same spectrum as example 4c, but annotated with the previously, incorrectly reported HLA peptide as described in Mylonas *et al*. Figure 2A. The non-matching synthetic peptide spectrum for the incorrect sequence is given as Box 1 example 4b.

## Box 1

Example use cases for Universal Spectrum Identifiers (USIs). See the [Example USIs] button at http://proteomecentral.proteomexchange.org/usi/ for a version of this box with active hyperlinks for exploring the spectra and PSMs.

### Case 1

Typical identification of an unmodified peptide in support of protein identification (one spectrum, simple interpretation, no mass modifications)

**mzspec:PXD000561:Adult_Frontalcortex_bRP_Elite_85_f09:scan:17555:VLHPLEGAVVIIFK/2**

*Example 1: Peptide Spectrum Match (PSM) of an unmodified doubly-charged peptide VLHPLEGAVVIIFK from the Kim et al*.20 *draft human proteome dataset. Most of the intense unannotated peaks are internal fragmentation ions of this peptide*.

### Case 2

Flexible notation for reporting identification of post-translational modifications (one spectrum, interpretation with Unimod names, Unimod identifiers, PSI-MOD names and PSI-MOD identifiers for mass modifications)

**mzspec:PXD000966:CPTAC_CompRef_00_iTRAQ_05_2Feb12_Cougar_11-10-09.mzML:scan:12298:[iTRAQ4plex]-HFFM[Oxidation]PGFAPLTSR/2**

*Example 2a: PSM of a iTRAQ4plex-labeled peptide from a CPTAC CompRef dataset*21, *with modifications specified using Unimod names*.

*Using names rather than accession numbers or mass deltas is the recommended notation since it precisely identifies the modification while also being easily interpretable*.

**mzspec:PXD000966:CPTAC_CompRef_00_iTRAQ_05_2Feb12_Cougar_11-10-09.mzML:scan:12298:[UNIMOD:214]-LHFFM[UNIMOD:35]PGFAPLTSR/2**

*Example 2b: Same CPTAC PSM with modifications specified using Unimod accession numbers. This notation is equally precise but not as easily readable as modification names*.

**mzspec:PXD000966:CPTAC_CompRef_00_iTRAQ_05_2Feb12_Cougar_11-10-09.mzML:scan:12298:[MOD:01499]-LHFFM[L-methionine sulfoxide]PGFAPLTSR/2**

*Example 2c: Same CPTAC PSM with modifications specified using PSI-MOD names and accession numbers*.

**mzspec:PXD000966:CPTAC_CompRef_00_iTRAQ_05_2Feb12_Cougar_11-10-09.mzML:scan:12298:[+144.1021]-LHFFM[+15.9949]PGFAPLTSR/2**

*Example 2d: Same CPTAC PSM with modifications specified using mass offsets. This notation is generally discouraged when the type of modification is known in advance, but is the only available option to report results of open modification searches returning algorithmically-detected uninterpreted mass offsets*.

### Case 3

Supporting evidence of translated gene products (i.e., protein existence) as detected in public datasets, including matches to spectra of synthetic peptides as required by the HUPO Human Proteome Project (HPP) guidelines for detection of novel proteins

**mzspec:PXD022531:j12541_C5orf38:scan:12368:VAATLEILTLK/2**

*Example 3a: Identification derived from a prey protein (Q5VTA0) in the Huttlin et al*.18 *BioPlex dataset, pulled down as a binding partner to bait protein C5orf38. With only this single identification, this protein remains an HPP missing protein since a single identification does not meet HPP guidelines*.

**mzspec:PXD022531:b11156_PRAMEF17:scan:22140:VAATLEILTLK/2**

*Example 3b: PSM of the same peptide as above, but derived from a recombinant protein used as a bait in the Huttlin et al. BioPlex dataset. This PSM provides a much higher signal-to-noise ratio synthetic peptide reference spectrum as required by HPP guidelines*.

### Case 4

Data reanalysis refuting previous claims of novel HLA peptides

**mzspec:PXD000394:20130504_EXQ3_MiBa_SA_Fib-2:scan:4234:SGVSRKPAPG/2**

*Example 4a: Identification originally used to reported a novel HLA peptide (Mylonas et al*.4 *Figure 2A)*.

**mzspec:PXD010793:20170817_QEh1_LC1_HuPa_SplicingPep_10pmol_G2_R01:scan:8296:SGVSRKPAPG/2**

*Example 4b: Spectrum of a synthetic peptide for the same peptide SGVSRKPAPG/2 revealing a distinctively different fragmentation pattern*

*from example 4a*.

**mzspec:PXD000394:20130504_EXQ3_MiBa_SA_Fib-2:scan:4234:ATASPPRQK/2**

*Example 4c: The same spectrum as for example 4a, but correctly identified to commonly-occurring peptide from UniProtKB protein Q9UQ35 (Mylonas et al. Figure 2B)*.

**mzspec:PXD010793:20170817_QEh1_LC1_HuPa_SplicingPep_10pmol_G2_R01:scan:7452:ATASPPRQK/2**

*Example 4d: Spectrum of a synthetic peptide for ATASPPRQK/2 (same as 4c) with a fragmentation pattern matching example 4c, thus*

*confirming the identification to protein Q9UQ35*.

### Case 5

Reporting spectra of unidentified peptides with the potential to lead to interesting new discoveries

**mzspec:PXD010154:01284_E04_P013188_B00_N29_R1.mzML:scan:31291**

*Example 5a: Unidentified peptide detected by clustering as highly abundant only in Small intestine and Duodenum out of 29 human tissues in PXD010154*.

**mzspec:PXD010154:01284_E04_P013188_B00_N29_R1.mzML:scan:31291:DQNGTWEM[Oxidation]ESNENFEGYM[Oxidation]K/2**

*Example 5b: Manual annotation of the spectrum from example 5a reveals it to be a multiply-modified version of a peptide previously detected*

*only as unmodified*.

The Human Proteome Project^16^ (HPP) has set a high bar for data quality and evidence in support of its goal to provide high-stringency detections for all human proteins. The latest version of its MS data interpretation guidelines 3.0^17^ have set a requirement that key detection claims of proteins not previously seen via MS must be accompanied by USIs referencing the key spectra for each claim, so that the peptide-spectrum matches can be transparently inspected by the community to verify their veracity. For example, the BioPlex dataset^18^ was important for detecting novel proteins that had not been previously observed^19^ but it was crucial to consider the provenance of every single identification to exclude all files from experiments where the protein was intentionally overexpressed (as per the standard protocol for analysis of protein-protein interactions). Example 3a in Box 1 provides a PSM derived from a prey protein pulled down as a binding partner to bait protein C5orf38. Example 3b provides a PSM of the same peptide as above, but derived from a recombinant protein used as a bait. This PSM provides a much higher signal-to-noise ratio synthetic peptide reference spectrum as required by HPP guidelines. Illustrating this application of USIs at a community-wide scale, MassIVE further provides an extensive list of USIs for 1,296,916 MassIVE-KB entries in support of HPP Protein Existence (PE) classifications for 16,393 proteins (available at http://massive.ucsd.edu/hpp), including USIs for matching spectra of synthetic peptides (when available in public datasets); an abridged version of this table is also provided as Supplementary Table 2.

## Conclusion

The Universal Spectrum Identifier provides a standardized mechanism for encoding a virtual path to any spectrum contained in data sets deposited to public repositories or contained in public spectral libraries. Current PX data repositories have implemented support for USIs as part of the submission or browsing process as described in the Supplementary Information. It has been primarily designed for the proteomics community, but could easily be adopted by the metabolomics community (and potentially others) if there is a will to do so (the current USI specification already supports non-proteomics spectra but additional guidelines would be needed for instance to represent non-proteomics identifications). It enables greater transparency and data ‘FAIRness’^2^ for providing spectral evidence in support of key findings in publications and public data, and provides a standardized mechanism for communicating identifiers of specific spectra among software applications. We thus expect that USIs will become a fundamental mechanism supporting open science across all fields of research supported by MS data.

## Notes

The authors declare no competing financial interest.

## Acknowledgements

This work was funded in part by the National Institutes of Health grants R01GM087221, R24GM127667, 1R01LM013115 and P41GM103484, and National Science Foundation grants 1933311, 1922871 and ABI 1759980. JAV wants to acknowledge Wellcome Trust grant number 208391/Z/17/Z, BBSRC BB/S01781X/1, BB/P024599/1 and the partnering grants BB/N022440/1 and BB/N022432/1. SK wants to acknowledge the Database Integration Coordination Program from the National Bioscience Database Center, Japan Science and Technology Agency [18063028] and the Japan Society for the Promotion of Science KAKENHI [JP20H03245]. We also would like to acknowledge the Research Foundation – Flanders (SB grant 1S90918N to TVDB; SB grant 1S50918N to RG; WB is a postdoctoral researcher).

## Notes

https://psidev.info/usi

### Competing Interest Statement

The authors have declared no competing interest.

### Summary of Updates

Minor additions as requested by reviewers. No substantial change.

