## Supplementary Information for "Universal Spectrum Identifier for mass spectra"

#### Supplementary Table 1

List of resources and their URLs that allow a user to copy-paste a USI into the web page and visualize the spectrum if it is available

| Resource | URL |
|---|---|
| ProteomeCentral | http://proteomecentral.proteomexchange.org/usi/ |
| jPOST | https://repository.jpostdb.org/spectrum/ |
| MassIVE | http://massive.ucsd.edu/ProteoSAFe/usi.jsp |
| PeptideAtlas | https://db.systemsbiology.net/sbeams/cgi/PeptideAtlas/ShowObservedSpectrum |
| PRIDE | https://www.ebi.ac.uk/pride/archive/spectra |
| ProteomicsDB | https://www.proteomicsdb.org/use/ |
| GNPS (metabolomics) | https://gnps.ucsd.edu/ProteoSAFe/gnps_usi.jsp |
| iProX | https://www.iprox.org/page/spectrum.html |

### USI Implementation notes from each resource

#### ProteomeCentral

ProteomeCentral hosts a web-based USI validator and resolver at http://proteomecentral.proteomexchange.org/usi/. Users of this web page may paste a USI into a text box and choose either to validate or fetch the spectrum/PSM encoded in the USI. Validation is performed by a back-end service that parses the supplied USI and decomposes it into its various components and ensures that they are well formed and follow the rules of the specification. Collection identifiers are checked to ensure that they encode a supported namespace, but not that they are within a valid range. MS runs components are separated but not checked as essentially any string is permitted. The indexType is checked for a valid value. The indexNumber is separated but not checked against the relevant file to ensure that it is within range. The interpretation string is parsed to ensure it is well formed and that controlled vocabulary (CV) terms are present in the relevant CVs. The service returns the decomposed and interpreted USI and a flag to indicate if it is valid or not. A well-formed USI may not actually correspond to a spectrum, but this cannot be determined without asking all resources to look up the USI.

The USI lookup function first validates the provided USI as described above, and then, if valid, sends the USI to all participating resource services that can provide spectra given a USI. USI lookups are performed in parallel within the JavaScript. A result table depicts the results from the participating resources as they arrive in real time. The first spectrum to return from one of the resources is sent to an embedded Lorikeet spectrum viewer in the page with basic defaults for an HCD MS/MS spectrum, along with the peptide interpretation, if provided. The user may click on various hyperlinks in the table to see the spectrum obtained from different resources, link out to the resources themselves, where the user can explore the spectrum using the native tools at the resource, as well as access additional information about the returned spectrum data. The USI lookup service may be triggered automatically without required copy-paste with a URL http://proteomecentral.proteomexchange.org/usi/?usi=<USI> where the <USI> is replaced with an actual USI, e.g. http://proteomecentral.proteomexchange.org/usi/?usi=mzspec:PXD000561:Adult_Frontalcortex_bRP_Elite_85_f09:scan:17555:VLHPLEGAVVIIFK/2.

#### jPOST

jPOST provides a USI resolver at https://repository.jpostdb.org/spectrum/. Currently there are 27M spectra corresponding to USIs. Spectra submitted by "Complete Submission" are basically available via USI. In addition, spectra submitted to PX partners other than jPOST will also provide USI from jPOST in the case of frequently used datasets.

#### MassIVE

The main MassIVE interface for resolution of USIs is available at https://massive.ucsd.edu/ProteoSAFe/usi.jsp. At the time of writing, MassIVE can resolve USIs for over >3.3 billion spectra in >10,000 datasets. In addition, MassIVE can also resolve USIs to >580 million Peptide Spectrum Matches (PSMs), including both PSMs that were included in complete dataset submissions (reported with PXD and MSV accessions) and PSMs derived by MassIVE (and third-party) reanalysis of public datasets (reported with RPXD and RMSV accessions). As illustrated in Figure S1, USIs can be entered in a text box and can be used to search MassIVE datasets in four different ways. Each of the four different types of search is accompanied by a small blue icon containing a short description of each type of search, as well as listing an example USI illustrating the types of results for each search.

The first option ("Search MassIVE spectra", also illustrated in Figure S1) is to search for the dataset and spectrum file(s) as specified in the USI mzspec:PXD000561:Adult_Frontalcortex_bRP_Elite_85_f09:scan:17555:VLHPLEGAVVIIFK/2. If successful, this search returns a table with one row per spectrum file with a matching file name. Clicking on the spectrum icon (on the left) for rows corresponding to spectrum files in standard open formats will open a new panel allowing for visualization and interactive exploration of the USI spectrum. If a peptide annotation is included in the USI then it is used to label and color spectrum peaks according to the theoretical masses of the standard fragment ion types.

The second option ("Search all MassIVE PSMs") is to search all of MassIVE for other possible identifications for the spectrum indicated in the USI. Since spectrum files can be resubmitted in multiple datasets and reanalyzed in multiple RPXD/RMSV reanalyses, this option searches for all occurrences of MassIVE PSMs matching the USI spectrum file and spectrum identifier (e.g., scan number). The example in Figure S2(a) shows that the initial spectrum identification (mzspec:PXD002041:TCGA-AA-3518-01A-11_W_VU_20120915_A0218_3F_R_FR04:scan:9649:ADNQRLKDENGALIR/2) reported in a proteogenomics search for a CPTAC dataset is actually a worse match to the spectrum than the alternative identification to a different peptide found in MassIVE reanalyses with a common N-term modification of +28 Da (likely Formylation, a common sample handling artefact), a type of modification that was not considered in the original paper.

The third option ("Search MassIVE-KB reference spectra") searches the MassIVE Knowledge Base (MassIVE-KB) for reference spectra for the same peptide as in the USI. Reference spectra can include spectra of synthetic peptides as well as spectra selected as representatives in the MassIVE-KB spectral library. As shown in Figure S2(b), the spectrum in the USI mzspec:PXD004452:20150410_QE3_UPLC9_DBJ_SA_46fractions_Rep1_18:scan:2873:PAPTLEEEKIR/2 was incorrectly identified to a peptide whose reference spectrum shows a distinct fragmentation pattern than that of the synthetic peptide reported in this search.

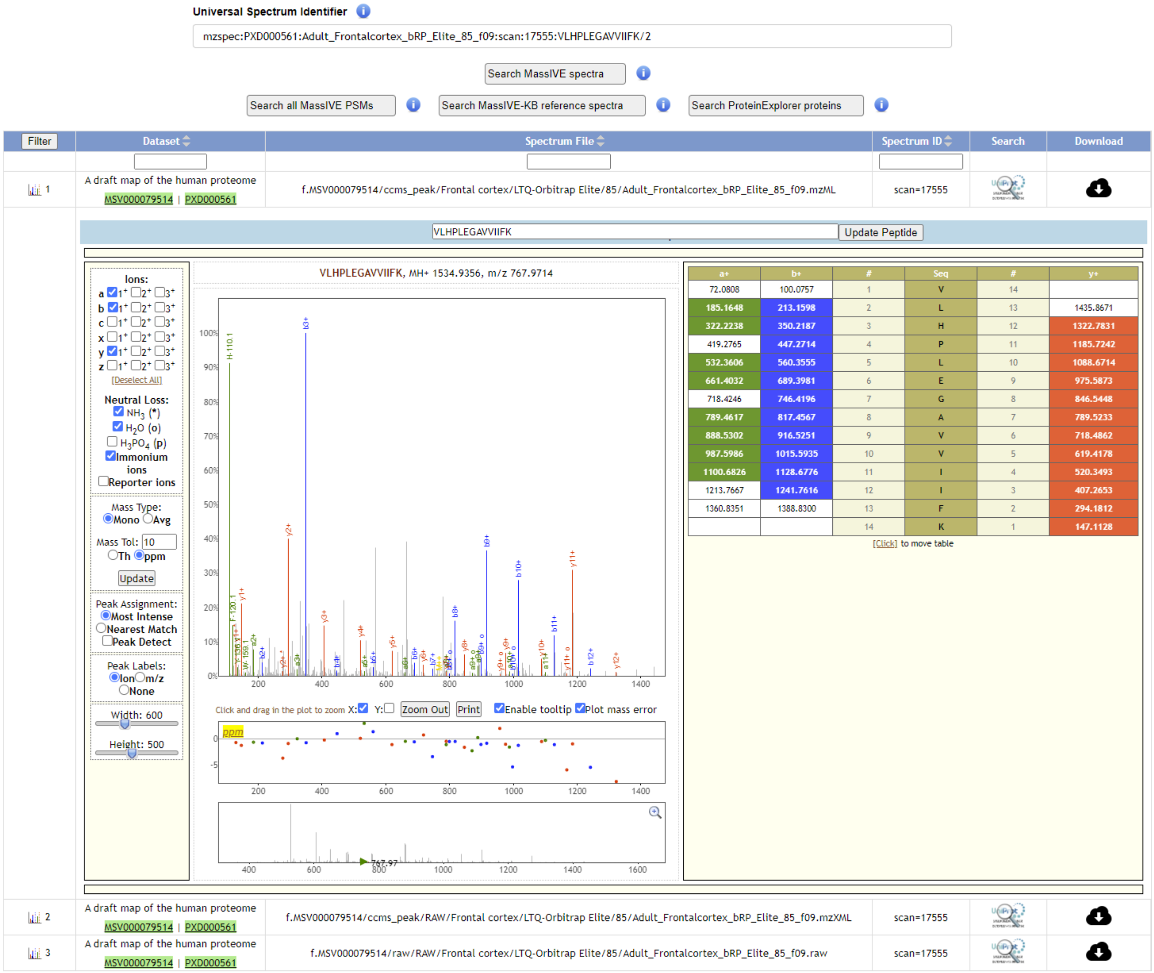


Figure S1. Main MassIVE functionality for resolution of Universal Spectrum Identifiers.

The fourth option ("Search Protein Explorer proteins") searches [MassIVE's Protein Explorer](#) for human proteins containing the USI peptide sequence. As shown in Figure S2(c), the peptide in the USI [mzspec:PXD000865:00644_H11_P004899_B0P_A00_R1:scan:3727:PAGDGTFQK/2](#) maps to 6 distinct HLA proteins.

To facilitate the generation of supplementary materials support the review process for the manuscripts associated with ProteomeXchange datasets, MassIVE also enables exporting identification results in TSV format including USIs for all selected results. This is described and illustrated in Figure S3.

(a) Searching for all reported identifications of the USI spectrum

mzspec:PXD000561:Adult_Frontalcortex_bRP_Elite_85_f09:scan:17555:VLHPLEGAVVIIFK/2

| Filter | Dataset | Spectrum File | Spectrum ID | Peptide |
|---|---|---|---|---|
| | | TCGA-AA-3518-C | scan=9649 | |
| 1 | CPTAC proteomic analysis of TCGA colon and rectal carcinomas using standard and customized databases, part 1 MSV000080147 \| PXD002041 | f.MSV000080147/peak/PEAK/TCGA-AA-3518-01A-11_W_VU_20120915_A0218_3F_R_FR04.mzML.gz | scan=9649 | ADNQRLKDENGALIR |
| 2 | MODa Re-analysis of CPTAC Patient 3518 RMSV000000233.2 | f.MSV000079852/peak/ColorectalCancer/TCGA-AA-3518-01A-11_Proteome_VU_20120915/mzML_data/TCGA-AA-3518-01A-11_W_VU_20120915_A0218_3F_R_FR04.mzML.gz | scan=9649 | A+28AVEEGIVLGGGC+57.021464ALLR |
| 3 | MODa Re-analysis of CPTAC Colorectal data RMSV000000004.4 \| RPXD006619.4 | f.MSV000079852/peak/ColorectalCancer/TCGA-AA-3518-01A-11_Proteome_VU_20120915/mzML_data/TCGA-AA-3518-01A-11_W_VU_20120915_A0218_3F_R_FR04.mzML.gz | scan=9649 | A+28AVEEGIVLGGGC+57.021464ALLR |
| 4 | MAESTRO Re-analysis of CPTAC Patient 3518 RMSV000000232.4 | f.MSV000080921/peak/TCGA-AA-3518-01A-11_Proteome_VU_20120915/mzML_data/TCGA-AA-3518-01A-11_W_VU_20120915_A0218_3F_R_FR04.mzML.gz | scan=9649 | A+28AVEEGIVLGGGC+57ALLR |
| 5 | Offical Proteogenomic CPTAC Search Results RMSV000000234.3 \| RPXD007509.3 | f.MSV000080147/peak/PEAK/TCGA-AA-3518-01A-11_W_VU_20120915_A0218_3F_R_FR04.mzML.gz | scan=9649 | ADNQRLKDENGALIR |

(b) Searching MassIVE-KB for reference spectra for the same peptide as in the USI

mzspec:PXD002041:TCGA-AA-3518-01A-11_W_VU_20120915_A0218_3F_R_FR04:scan:9649:ADNQRLKDENGALIR/2

| Filter | Library | Sequence | Peptide Length | #Matched Proteins | #Matched Proteins w/ 0-1 SAAV Mismatch | Unique Exon Match | Exon Junction Match |
|---|---|---|---|---|---|---|---|
| | | PAPTLEEEKIR | ~ | ~ | ~ | ~ | ~ |
| 1 | Bioplex Synthetics | PAPTLEEEKIR | 11 | 1 | 1 | 1 | 0 |

(c) Searching Protein Explorer for proteins containing USI peptide

mzspec:PXD000865:00644_H11_P004899_B0P_A00_R1:scan:3727:PAGDGTFQK/2

MassIVE Spectra Proteins Hits 1 - 6 out of 6 Go to Go Export Filtered Results

Select columns

| Apply Filters | Protein Accession | Gene | neXtprot Protein Existence | Total Unique Peptides | HUPO Peptides | Total Unique Exons | Total PSMs | Total Unique PSMs | Protein Description |
|---|---|---|---|---|---|---|---|---|---|
| Filter By: | | | ~ | ~ | ~ | ~ | ~ | ~ | |
| 1 | P01893 | HLA-H | 5 | 8 | 6 | 0 | 4122 | 283 | Putative HLA class I histocompatibility antigen, alpha chain H |
| 2 | P04439 | HLA-A | 1 | 79 | 22 | 0 | 7565 | 3683 | HLA class I histocompatibility antigen, A alpha chain |
| 3 | P10321 | HLA-C | 1 | 39 | 18 | 0 | 7753 | 2624 | HLA class I histocompatibility antigen, C alpha chain |
| 4 | P13747 | HLA-E | 1 | 81 | 30 | 0 | 4864 | 2425 | HLA class I histocompatibility antigen, alpha chain E |
| 5 | P17693 | HLA-G | 1 | 22 | 9 | 0 | 1592 | 416 | HLA class I histocompatibility antigen, alpha chain G |
| 6 | P30511 | HLA-F | 1 | 34 | 21 | 0 | 2175 | 1501 | HLA class I histocompatibility antigen, alpha chain F |

Figure S2. Advanced MassIVE functionality for resolution of Universal Spectrum Identifiers.

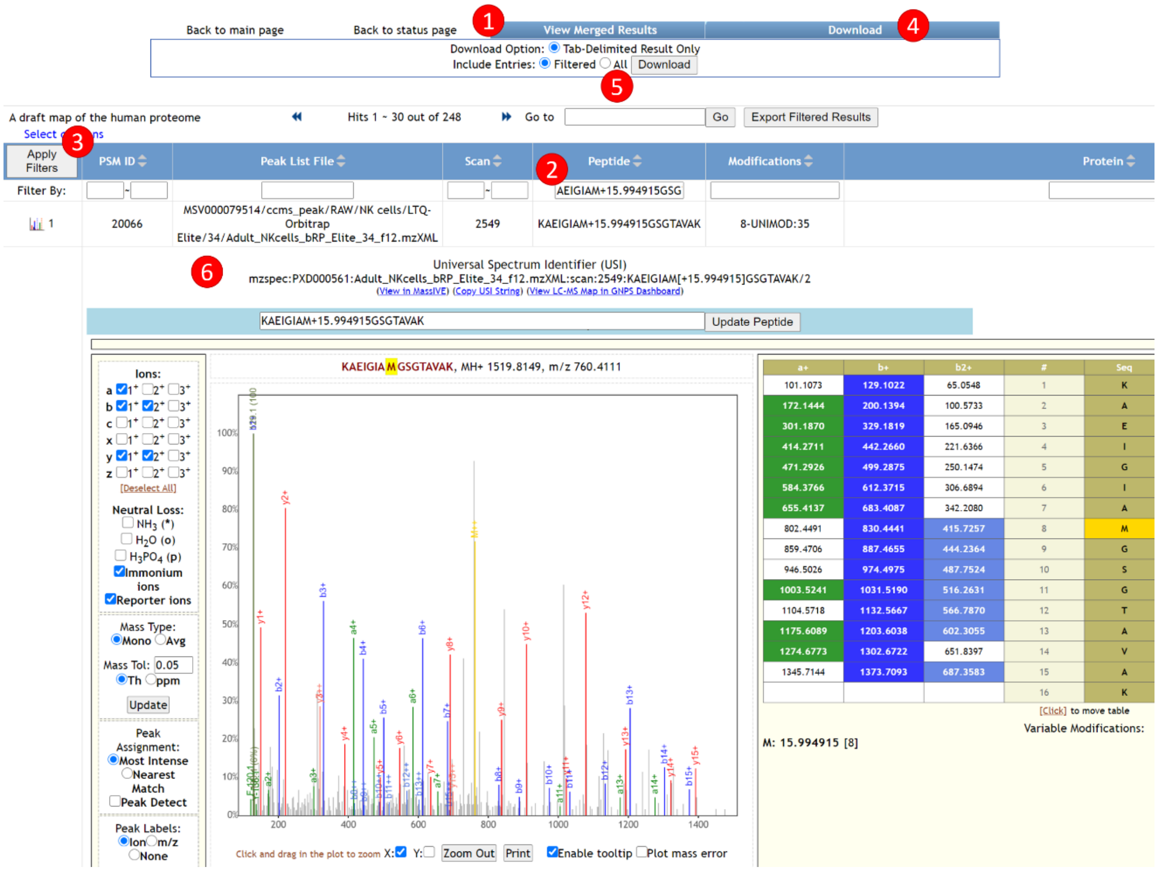


Figure S3. MassIVE functionality for exporting Universal Spectrum Identifiers for subsets of identified spectra, as is commonly important for identifications of modified peptides. This example illustrates how to export USIs for all identifications with an oxidized methionine on sequence AEIGIAMGSGTAVAK in one of the original human draft proteome datasets. Starting with "Browse Results" in the MassIVE dataset page for PXD000561, the figure illustrates that **(1)** clicking View Merged Results shows all identifications in the dataset, regardless of the number of separate searches; **(2)** entering a filter string in the Peptide field and **(3)** clicking Apply Filters selects all spectrum identifications matching the filter text; **(4)** clicking the Download tab and **(5)** selecting to download Filtered results produces a TSV file with all USIs for all filtered results that can be submitted with a manuscript to support the review process for the manuscript associated with a dataset. In addition, **(6)** USIs are also show for every row in the filtered results when clicking on the spectrum icon (leftmost column) to visualize the spectrum identification. The same results can also be communicated and accessed directly using the same URL that was used to generate this figure.

### PeptideAtlas

PeptideAtlas provides access to over 1 billion spectra via the USI mechanism. Users can go directly to https://db.systemsbiology.net/sbeams/cgi/PeptideAtlas/ShowObservedSpectrum and copy-paste a USI. If the dataset is housed at PeptideAtlas, then the system dynamically finds the location of the dataset on PeptideAtlas disks, finds the mzML.gz file corresponding to the requested MS run, and then extracts the requested spectrum from the mzML.gz file. Part of the PeptideAtlas processing pipeline is to download raw vendor files from the original hosting repositories and convert them to mzML.gz via msconvert or ThermoRawFileReader and these are all stored in an accessible manner. The extracted spectrum is then displayed with the requested annotation embedded in the USI if there is one.

Additionally, as users are browsing the PeptideAtlas web interface, if they click on a link to view a PSM in a PeptideAtlas build, if the dataset has a PXD identifier (not all datasets in PeptideAtlas do), then a USI for that PSM is created and displayed above the spectrum.

#### PRIDE

At the time of writing, the PRIDE database provides access to over 540 million peptide-spectrum matches (PSMs) originally submitted by researchers in the PRIDE Archive Spectra resource (https://www.ebi.ac.uk/pride/archive/spectra). Users can search by USIs, which enables them to find specific PSMs from submitted data.

In addition, a peptide sequence search can be performed in the search-box, which enables users to find PSMs across the entire resource. A list of PSMs is shown (Figure S4) including peptide sequences, post-translational modifications, search engine scores, charge, and two additional columns that highlight whether the PSM has passed or not the original analysis threshold and PRIDE internal pipelines thresholds (e.g. PSM FDR < 0.1).

The accession column in the result table provides a direct link to the project centric page which contains the information of the original submitted dataset, including the results details page. If users perform a search using a USI (e.g. https://www.ebi.ac.uk/pride/archive/spectra?usi=mzspec:PXD000966:CPTAC_CompRef_00_iTRAQ_12_5Feb12_Cougar_11-10-11.mzML:scan:11850:[UNIMOD:214]YYWGGLYSWDMSK[UNIMOD:214]/2), the corresponding spectra will be shown in the spectrum panel below in the same page.

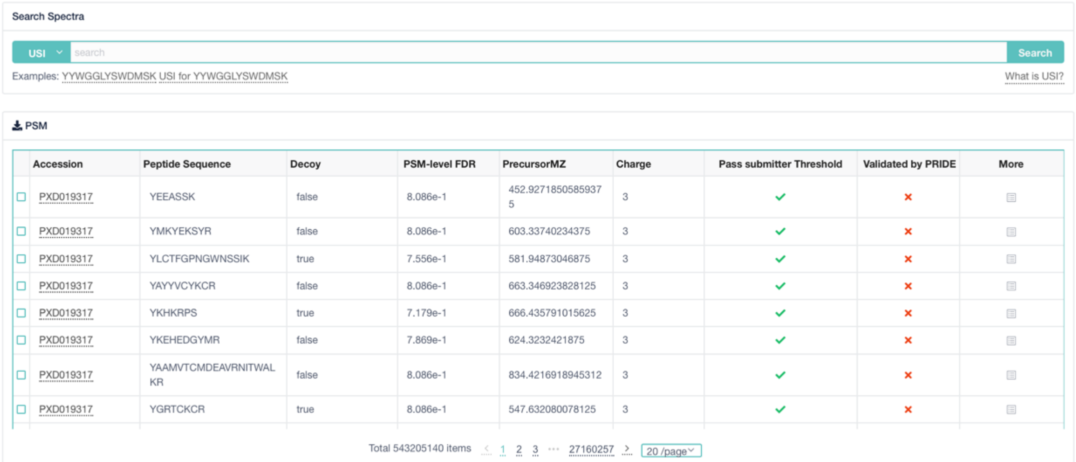


| Accession | Peptide Sequence | Decoy | PSM-level FDR | PrecursorMZ | Charge | Pass submitter Threshold | Validated by PRIDE | More |
|---|---|---|---|---|---|---|---|---|
| PXD019317 | YEEASSK | false | 8.086e-1 | 452.92718505859375 | 3 | ✓ | × | |
| PXD019317 | YMKYEKSYR | false | 8.086e-1 | 603.33740234375 | 3 | ✓ | × | |
| PXD019317 | YLCTFGPNGWNSSIK | true | 7.556e-1 | 581.94873046875 | 3 | ✓ | × | |
| PXD019317 | YAYYVCYKCR | false | 8.086e-1 | 663.346923828125 | 3 | ✓ | × | |
| PXD019317 | YKHKRPS | true | 7.179e-1 | 666.435791015625 | 3 | ✓ | × | |
| PXD019317 | YKEHEDGYMR | false | 7.869e-1 | 624.3232421875 | 3 | ✓ | × | |
| PXD019317 | YAAMVTCMDEAVRNITWALKR | false | 8.086e-1 | 834.4216918945312 | 3 | ✓ | × | |
| PXD019317 | YGRTCKCR | true | 8.086e-1 | 547.632080078125 | 3 | ✓ | × | |



Figure S4. PRIDE Archive Spectra (https://www.ebi.ac.uk/pride/archive/spectra) provides access to millions of PSMs.

It is possible to access and visualise any PSM included in a publicly available "complete" PRIDE dataset. To do that, users need to go to https://www.ebi.ac.uk/pride/archive/spectra and search for a particular peptide sequence.

For instance, for the peptide sequence WQLVGITSWGEGCAR, the resulting URL is: https://www.ebi.ac.uk/pride/archive/spectra?peptideSequence=WQLVGITSWGEGCAR (Figure S5). At that point it is possible to select all PSMs containing that particular sequence in a given dataset (Figure S6).

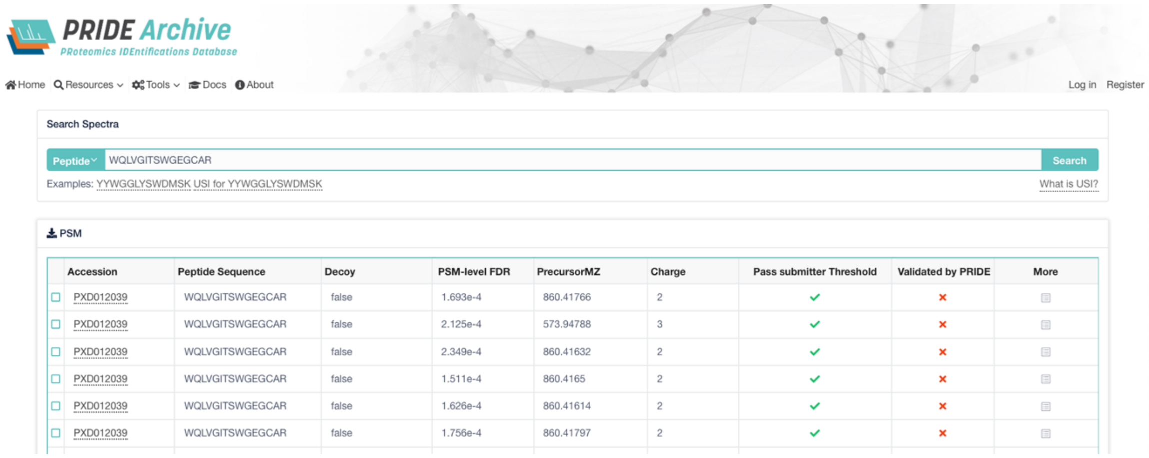


| Accession | Peptide Sequence | Decoy | PSM-level FDR | PrecursorMZ | Charge | Pass submitter Threshold | Validated by PRIDE | More |
|---|---|---|---|---|---|---|---|---|
| PXD012039 | WQLVGITSWGEGCAR | false | 1.693e-4 | 860.41766 | 2 | ✓ | × | |
| PXD012039 | WQLVGITSWGEGCAR | false | 2.125e-4 | 573.94788 | 3 | ✓ | × | |
| PXD012039 | WQLVGITSWGEGCAR | false | 2.349e-4 | 860.41632 | 2 | ✓ | × | |
| PXD012039 | WQLVGITSWGEGCAR | false | 1.511e-4 | 860.4165 | 2 | ✓ | × | |
| PXD012039 | WQLVGITSWGEGCAR | false | 1.626e-4 | 860.41614 | 2 | ✓ | × | |
| PXD012039 | WQLVGITSWGEGCAR | false | 1.756e-4 | 860.41797 | 2 | ✓ | × | |

Figure S5. Screenshot showing the result from searching for the peptide sequence WQLVGITSWGEGCAR.

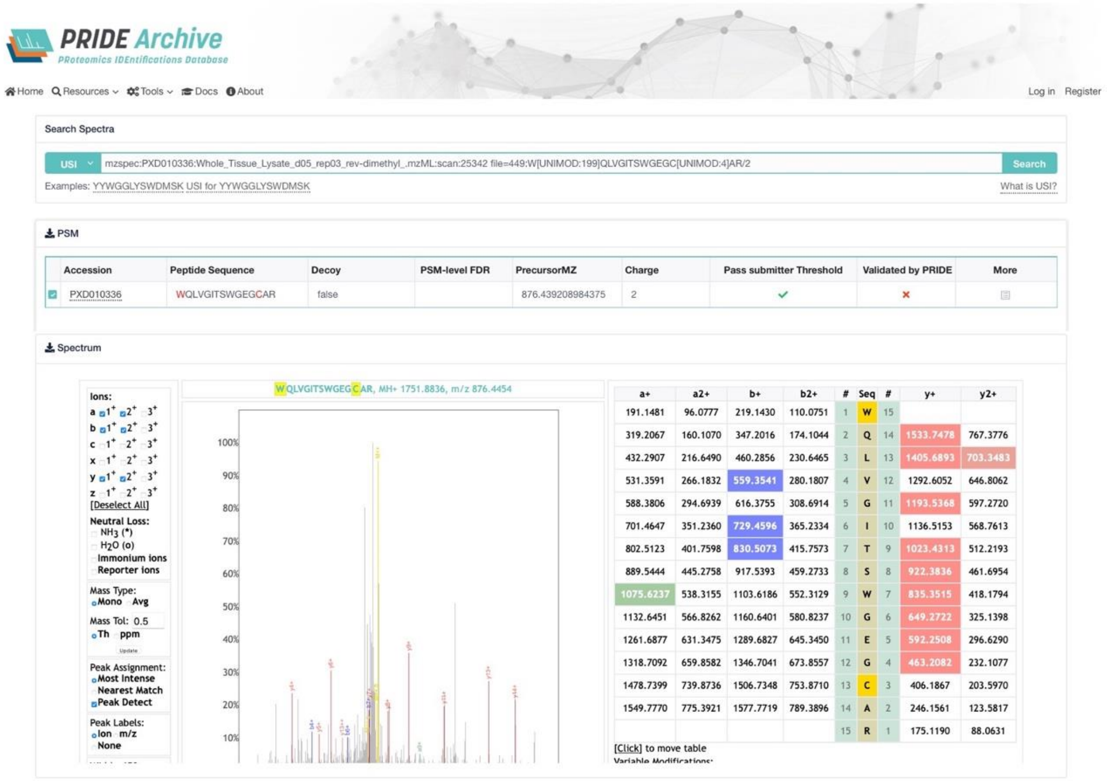


Figure S6. Screenshot showing the mass spectrum corresponding to the selected PSM (peptide sequence WQLVGITSWGEGCAR in dataset PXD010336).

#### ProteomicsDB (Universal Spectrum Explorer)

The ProteomicsDB team has developed a web tool called Universal Spectrum Explorer (USE - https://www.proteomicsdb.org/use/)[DOI:10.1101/2020.09.08.287557v1]; to query and visualize public spectra using USIs from multiple archives including PRIDE and PeptideAtlas. In addition, USE can retrieve real-time prediction of tandem mass spectra from the deep learning framework Prosit. The query spectra from public USI or provided by the user can be exported from USE as editable scalable high quality vector graphics (e.g., https://www.proteomicsdb.org/use/?usi=mzspec:PXD015890:18May18_Olson_WT2.raw%20(F001551).mzid_18May18_Olson_WT2.raw_(F001551).MGF:index:6913:AEAEAQAEELSFPR/2&usi_origin=pride).

#### GNPS (metabolomics)

GNPS hosts the Metabolomics Spectrum Resolver tool at https://gnps.ucsd.edu/ProteoSAFe/gnps_usi.jsp [DOI:10.1101/2020.05.09.086066] to access metabolomics mass spectra available in MassIVE/GNPS and alternative metabolomics data repositories. Besides spectra deposited to GNPS [DOI:10.1038/nbt.3597], the Metabolomics Spectrum Resolver builds upon the USI standard to resolve spectra from other popular metabolomics data resources, such as MetaboLights [DOI:10.1093/nar/gks1004], Metabolomics Workbench [DOI:10.1093/nar/gkv1042], MassBank [DOI:10.1002/jms.1777], MoNA, and MS2LDA [DOI:10.1093/bioinformatics/btx582] to provide a universal interface to more than 450 million metabolomics spectra. The Metabolomics Spectrum Resolver can display individual spectra or display two spectra in a mirror plot to evaluate their similarity. Additionally, it supports linking back to the source spectra in their original data resources to allow users to explore the original context of the spectra and computes SPLASH identifiers [DOI:10.1038/nbt.3689] to enable comparisons of spectra across different resources.

#### iProX

iProX provides USI search and display at http://www.iprox.org/page/spectrum.html for 6 million spectra in the HBase. The result file of the "complete submission" project in iProX are parsed into spectra records uniquely marked by USIs and stored in the HBase. There is also a backup mechanism to ensure data reliability.
